# Tissue-specific overexpression of the double-stranded RNA transporter SID-1 limits lifespan in *C. elegans*

**DOI:** 10.1101/2022.11.23.517635

**Authors:** Henrique Camara, Carlos A. Vergani-Junior, Silas Pinto, Thiago L. Knittel, Willian G. Salgueiro, Guilherme Tonon-da-Silva, Juliana Ramirez, Evandro A. De-Souza, Marcelo A. Mori

## Abstract

Intertissue RNA transport has emerged as a novel signaling mechanism. In *C. elegans*, this is conferred by the systemic RNAi pathway, in which the limiting step is the cellular import of extracellular RNAs via SID-1. To better understand the physiological role of systemic RNAi *in vivo*, we modified the function of SID-1 through loss-of-function mutation and tissue-specific overexpression of *sid-1* in *C. elegans*. We observed that *sid-1* loss-of-function mutants are as healthy as wild-type worms. Conversely, overexpression of *sid-1* in intestine, muscle, or neurons rendered worms short-lived. The effects of intestinal *sid-1* overexpression were reversed by silencing the components of the systemic RNAi pathway *sid-1, sid-2* and *sid-5*, thus implicating RNA transport. Moreover, silencing the miRNA biogenesis proteins *pash-1* and *dcr-1* rendered the lifespan of worms with intestinal *sid-1* overexpression similar to controls. Lastly, we observed that the lifespan decrease produced by tissue-specific *sid-1* overexpression was dependent on the bacterial food source. Collectively, our data support the notion that systemic RNA signaling is tightly regulated, and unbalancing that process provokes a reduction in lifespan.

## INTRODUCTION

Non-coding RNAs are found in extracellular fluids coupled with proteins or within vesicles such as exosomes (Turchinovich *et al*., 2013). Specific extracellular non-coding RNAs (ncRNAs) have been shown to control gene expression cell non-autonomously, thus contributing to whole-body metabolic homeostasis, from nematodes to mammals (Mori *et al*., 2019; Zhou *et al*., 2019). As with hormones, extracellular ncRNA signaling is expected to be regulated at the level of secretion, extracellular transport, and recognition by the target cell. In mammals, important advances have been made in the characterization of RNA secretion mechanisms. Cell-type-specific motifs determine the secretion or intracellular retention of miRNAs (Garcia-Martin *et al*., 2022). These motifs are likely recognized by a long list of sorters that direct the RNAs for specific subcellular components, such as the late endosomes for secretion or the nucleus for cellular retention (Ruiz *et al*., 2021). However, specific effectors of mammalian extracellular ncRNA signaling are largely unknown, which poses a barrier to selectively manipulating this pathway.

Conversely, a well-described pathway for intertissue RNA transport is present in the nematode *C. elegans*. This pathway is referred to as the systemic RNAi pathway, which distributes endogenous and exogenous double-stranded RNAs (dsRNAs) throughout the body and elicits gene silencing in all cells of *C. elegans* except neurons, where this pathway is less active due to low expression of the dsRNA transporter SID-1 (Winston, Molodowitch and Hunter, 2002; Calixto *et al*., 2010). SID-1 is a conserved transmembrane channel operated by dsRNA which allows selective transport of RNAs once activated (Feinberg and Hunter, 2003; Shih and Hunter, 2011). The transport of RNAs via SID-1 is passive and has no size restriction or sequence specificity, but requires an RNA:RNA duplex structure (double-stranded structured single-stranded RNAs, such as RNA hairpins and pre-miRNAs are also transported) (Shih *et al*., 2009). Importantly, SID-1 expression is sufficient to sensitize neuronal cells of *C. elegans* (Calixto et al. 2010) and *Drosophila* S2 cells (Shih and Hunter, 2011) to extracellular dsRNAs. Additionally, neuronal expression of SID-1 reduces systemic RNAi efficiency in other somatic cells of the worm, suggesting that the distribution of extracellular RNAs is controlled at the level of SID-1 expression (Calixto *et al*., 2010).

With this in mind, we manipulated *sid-1* expression by loss-of-function mutations and tissuespecific overexpression as models of altered extracellular RNA transport and assessed the physiological consequences of these manipulations. We observed that *sid-1* mutants were as healthy as wild-type worms in a variety of phenotypic assays. In contrast, tissue-specific *sid-1* overexpression in the intestine, muscle, or neurons rendered worms short-lived, a phenotype dependent on components of the systemic RNAi and the miRNA pathways. Additionally, lifespan reduction was influenced by the bacterial food source and partially rescued by the downregulation of the environmental RNA transporter SID-2. Collectively, these data demonstrate the importance of a properly balanced intertissue distribution of extracellular non-coding RNAs to determine longevity in *C. elegans*.

## RESULTS & DISCUSSION

### *Sid-1* mutants are as healthy as wild-type worms

To investigate if dsRNA import plays a role in *C. elegans* physiology, we performed a phenotypic screen in three different loss-of-function mutant alleles of *sid-1* (*qt2, qt9*, and *pk3321*). As general physiological parameters, we measured brood size, developmental time, and lifespan. We did not observe differences between the *sid-1* mutants and the wild-type worm (Figure 1A-D). To check whether SID-1 is necessary for stress response, we measured the survival of *sid-1* mutants under acute oxidative stress or chronic thermic stress conditions. Survival of *sid-1* mutants was similar to the controls in both protocols (Figure 1E, F). Thus, *sid-1* loss-of-function does not impair *C. elegans* longevity, reproductive capacity, development or stress response.

**Figure 1.**
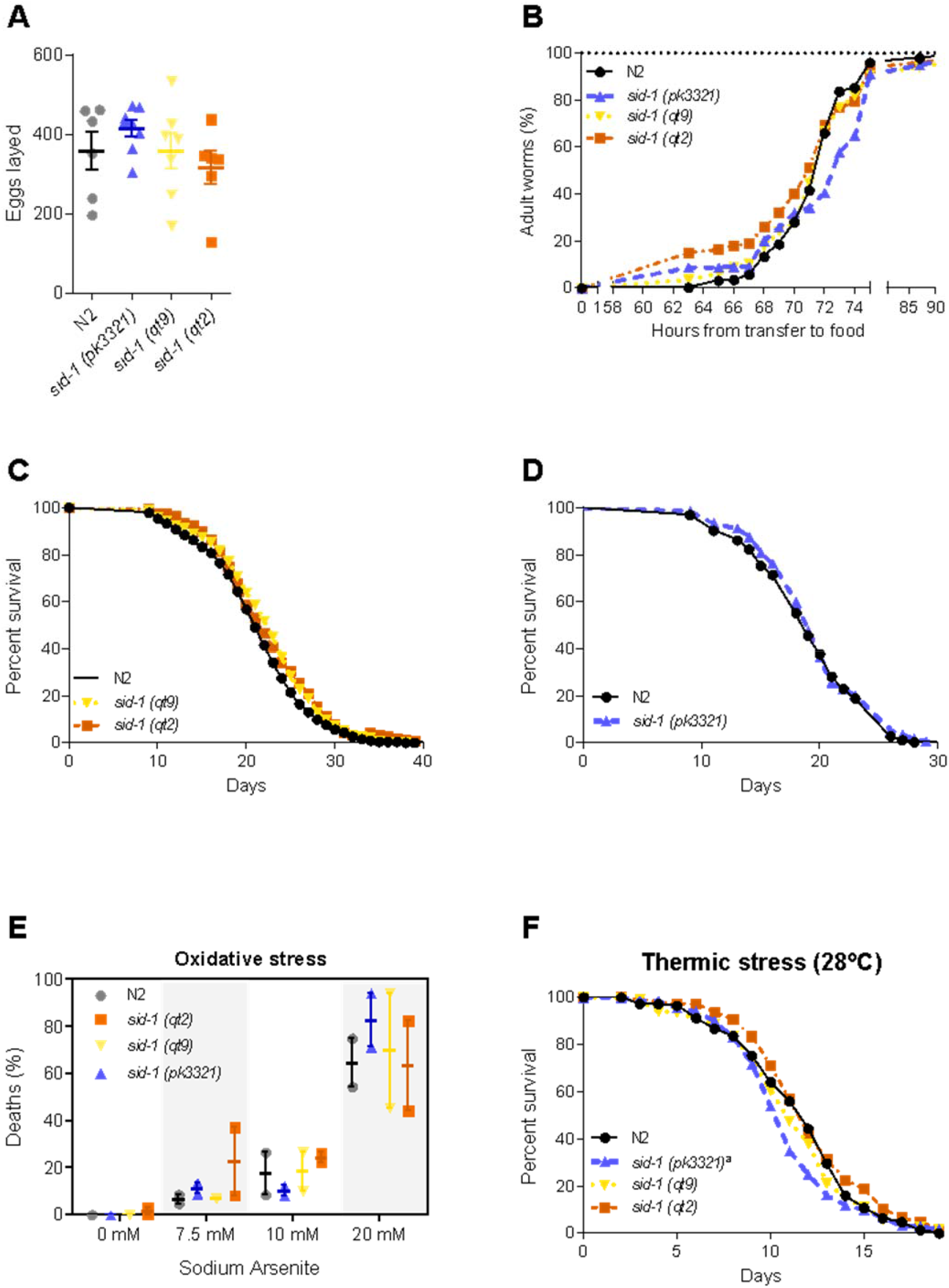
Phenotypic characterization of *sid-1* mutants. (A) Total egg laying of wild-type (N2) and *sid-1* mutants. Data are expressed as mean ± S.E.M (n = 6-8, One-way ANOVA with Dunnett’s multiple comparison test). (B) Developmental time of synchronized wild-type and *sid-1* mutants measured as time to reach adulthood. Data is the average of three independent trials [N2, n = 340; *sid-1* (*pk3321*), n = 231; *sid-1* (*qt9*), n = 269; *sid-1* (*qt2*), n = 212] and compared by the log-rank test. (C, D) Lifespan of control (N2) or *sid-1* mutants maintained at 20°C. (E) Percentage of alive wild-type and *sid-1* mutant worms after six to eight hours of exposure to 7.5 mM, 10 mM or 20 mM sodium arsenite. Data are expressed as mean ± S.E.M (n= 30-90, Two-way ANOVA, P<0.05 for arsenite concentration effect). (F) Lifespan of control (N2) or *sid-1* mutants maintained at 28°C. In C, D, and F, lifespan was compared by the log-rank test (^a^P<0.05 in comparison to wild-type control). Data are the average of two to three independent trials. See Supplementary Table 1 for lifespan statistics.

### Tissue-specific SID-1 overexpression decreases lifespan

Cell-type specific overexpression of *sid-1* is sufficient to increase dsRNA transport (Feinberg and Hunter, 2003; Shih *et al*., 2009; Calixto *et al*., 2010; Shih and Hunter, 2011). Therefore, we analyzed transgenic worms with tissue-specific overexpression of SID-1 in the intestine, muscle, or neurons as models of exacerbated import of extracellular dsRNAs into these selected tissues. We also crossed these transgenic worms with *sid-1 (qt9*) mutants to generate tissue-specific *sid-1* rescue strains. Overexpression of *sid-1* was confirmed in all the models (Supplementary Figure 1).

**Supplementary Figure 1.**
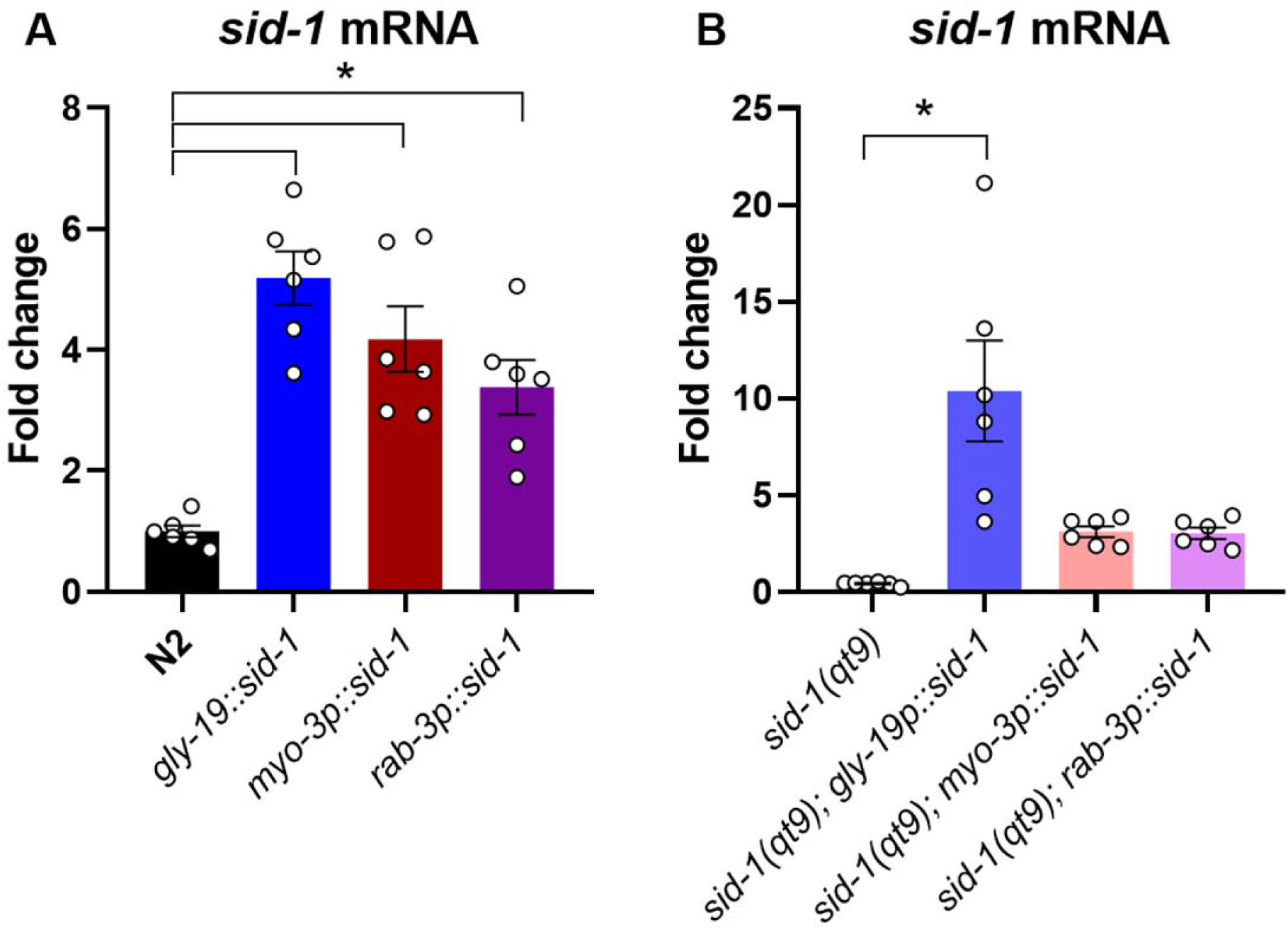
Validation of *sid-1* overexpression. Levels of *sid-1* on D0 adult control (N2) (A) or *sid-1* (*qt9*) mutants (B) with intestinal (*gly-19p::sid-1*), muscular (*myo-3p::sid-1*), and neuronal (*rab-3p::sid-1*) transgenic overexpression of *sid-1* (*P<0.05 vs. N2, n=6, One-way ANOVA, Dunnett’s multiple comparison test).

Intestinal overexpression of *sid-1* had the greatest impact on longevity and reduced the median lifespan of worms on the wild-type or *sid-1 (qt9*) background by 24.0 ± 1.5% and 12.8 ± 3.5%, respectively (Figure 2A). Muscle overexpression of *sid-1* also reduced the median lifespan of worms on the wild-type or *sid-1 (qt9*) backgrounds, although to a lesser magnitude (8.5 ± 4.1% and 7.0 ± 5.4%, respectively) (Figure 2B). Lastly, the neuronal overexpression of *sid-1* reduced the median lifespan of worms on the wild-type or *sid-1 (qt9*) background by 7.0 ± 5.4% and 14.7 ± 6%, respectively (Figure 2C). Thus, in contrast to the intestine and muscle *sid-1* overexpression models, the neuronal *sid-1* overexpression was worse in worms carrying the *sid-1 (qt9*) mutant allele than in the wild-type *sid-1* background. These data indicate that tissue-specific increase of SID-1 reduces the lifespan of *C. elegans*.

**Figure 2.**
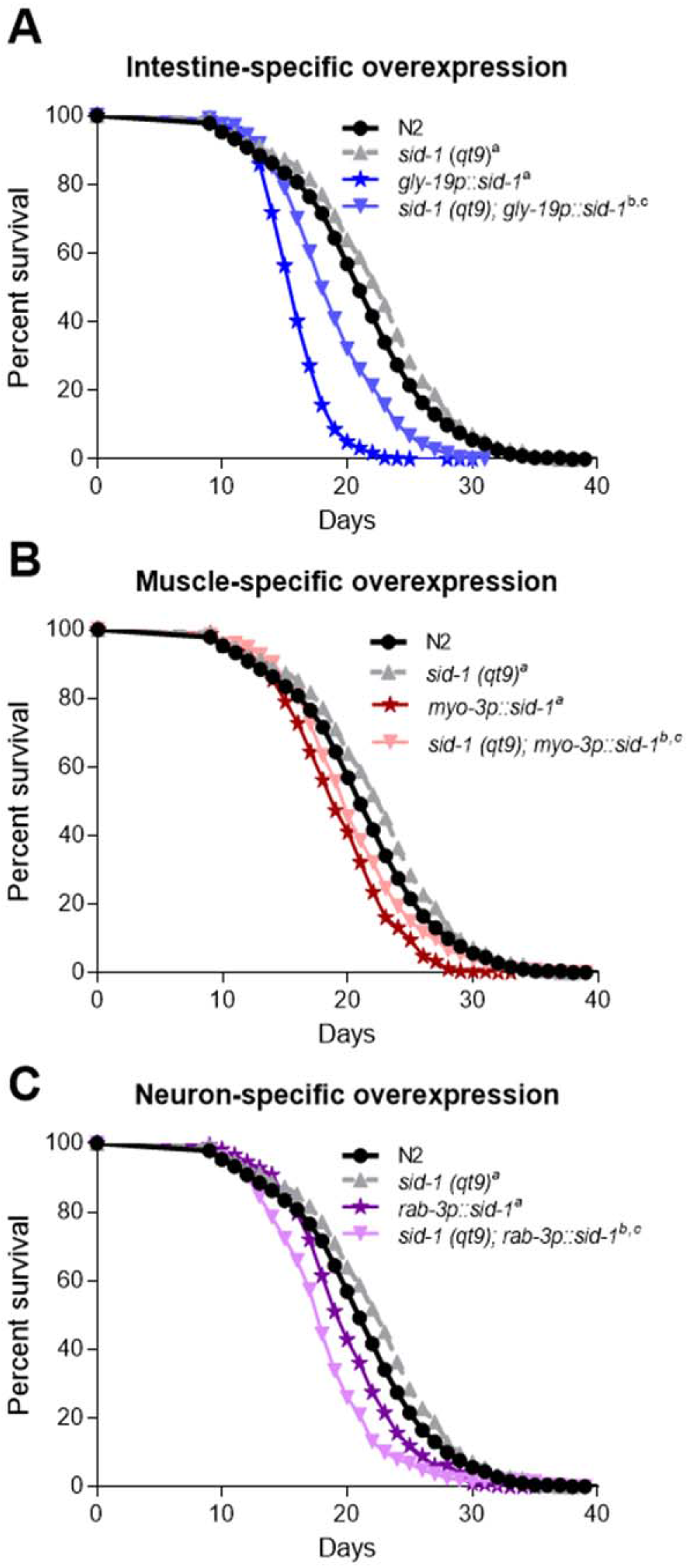
Tissue-specific increase of SID-1 decreases lifespan. Lifespan at 20°C of controls (N2) or *sid-1* (*qt9*) mutants with intestinal (*gly-19p::sid-1* [A]), muscular (*myo-3p::sid-1* [B]), and neuronal (*rab-3p::sid-1* [C]) *sid-1* overexpression. Lifespan was compared by the log-rank test (^a^P<0.05 in comparison to wild-type control; ^b^P<0.05 in comparison to *sid-1(qt9);* ^c^P<0.05 in comparison to *sid-1* overexpression). Data are the average of three independent trials. See Supplementary Table 1 for lifespan statistics.

### Tissue-specific SID-1 overexpression interacts with the systemic RNAi and miRNA pathways to control lifespan

The canonical pathway of environmental systemic RNAi begins with the uptake of dsRNA by the intestinal cells via SID-2 (Winston *et al*., 2007). SID-5 is then required to spread functional RNAs to distant cells (Hinas, Wright and Hunter, 2012), where they are taken up by SID-1 to produce gene silencing (Jose *et al*., 2012). To test whether the lifespan reduction induced by SID-1 overexpression interacted with the systemic RNAi pathway, we silenced *sid-1, sid-2* or *sid-5 in the* intestinal *sid-1 over*expression model, as this had the greatest lifespan reduction in our experiments. Consistent with the longevity of *sid-1* mutants, RNAi targeting *sid-1* did not affect the lifespan of wild-type worms. Conversely, *sid-1* RNAi restored the lifespan of worms overexpressing *sid-1* in the intestine to wild-type levels, validating that SID-1 overexpression mediates the observed lifespan reduction (Figure 3A). Similarly, RNAi targeting the RNA transporter *sid-5* or the environmental RNA importer *sid-2* partially reversed the lifespan reduction produced by intestinal *sid-1* overexpression (Figure 3B), reinforcing the role of systemic RNA transport in the phenotype and pointing towards a potential effect of exogenous RNA.

**Figure 3.**
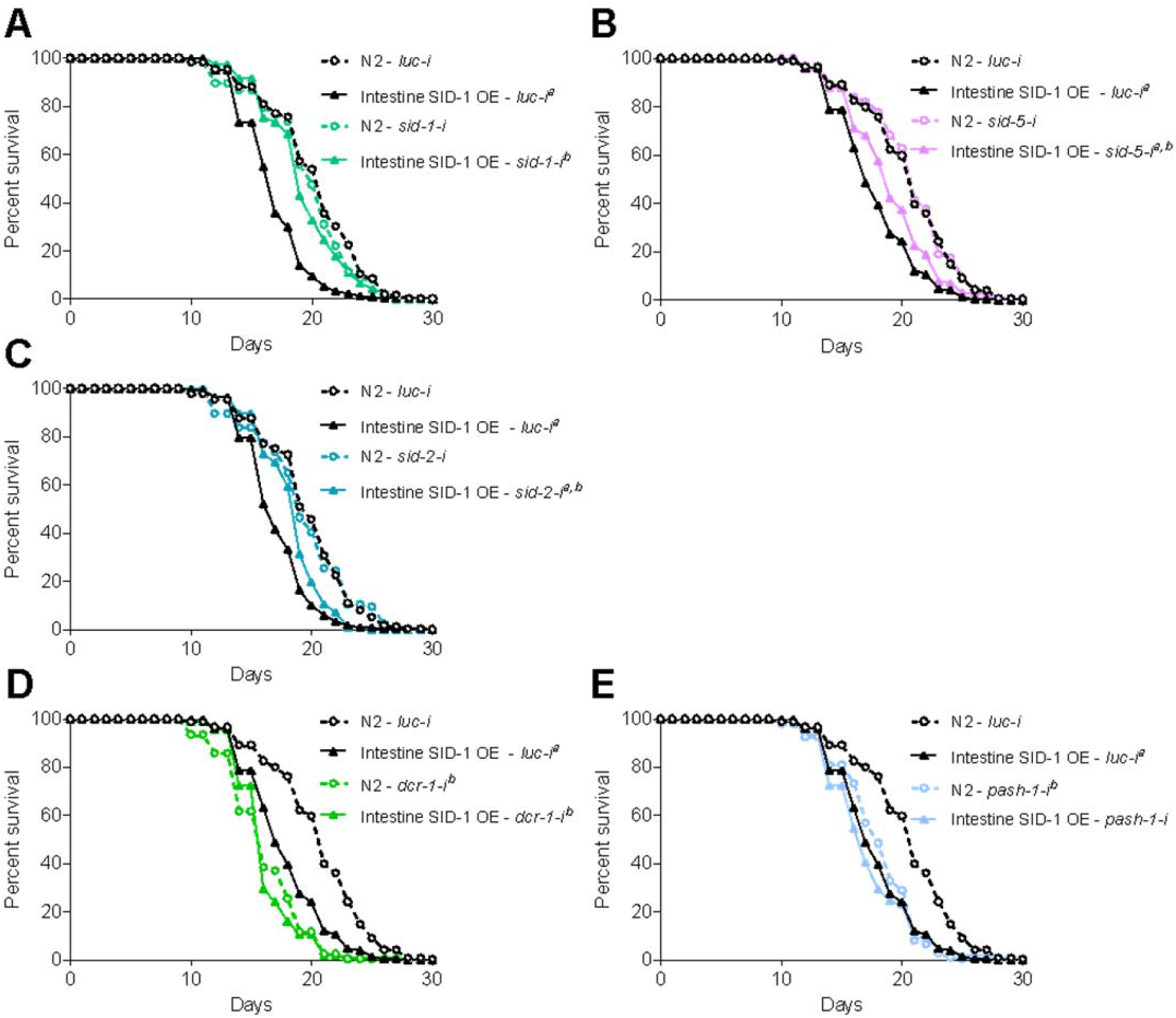
Lifespan reduction of tissue-specific *sid-1* expression interacts with the systemic RNAi and miRNA pathways. Lifespan of control (N2) and intestinal *sid-1* overexpressing (*gly-19p::sid-1*) animals maintained at 20°C and fed the RNAi targeting luciferase or *sid-1* (A), *sid-5* (B), *sid-2* (C), *dcr-1* (D) and *pash-1* (E). Lifespan was compared by the log-rank test (^a^P<0.05 in comparison to wild-type control; ^b^P<0.05 in comparison to luciferase RNAi control). Data are the average of two (A, C) and three (B, D, E) independent trials. See Supplementary Table 1 for lifespan statistics.

SID-1 transports a wide variety of dsRNA structures, including long or short dsRNAs and double-stranded structured ssRNA (e.g., pre-miRNAs) (Shih and Hunter, 2011). To unravel the potential RNA species involved in lifespan reduction induced by tissue-specific *sid-1* expression, we performed an RNAi screen for key components of the endo-siRNA (*dcr-1, ergo-1, hrde-1*, and *nrde-3*), exo-siRNA (*dcr-1* and *rde-4*), miRNA (*dcr-1* and *pash-1*), piRNA (*csr-1, unc-130* and *prg-1*), and antiviral pathways (*drh-1*). The list of RNAi clones tested is presented in Supplementary Table 2.

RNAi targeting components of the miRNA biogenesis pathway (*i.e., dcr-1* and *pash-1*) rendered the lifespan of worms with intestinal *sid-1* transgenic expression indistinguishable from their wild-type controls (Figure 3D-E). RNAi targeting other small RNA pathway genes did not affect the lifespan difference between the wild-type and the strain overexpressing *sid-1* in the intestine (Supp. Figure 2). These results demonstrate that systemic RNA transport and miRNAs are necessary for intestinal *sid-1* overexpression to reduce *C. elegans* lifespan.

**Supplementary Figure 2.**
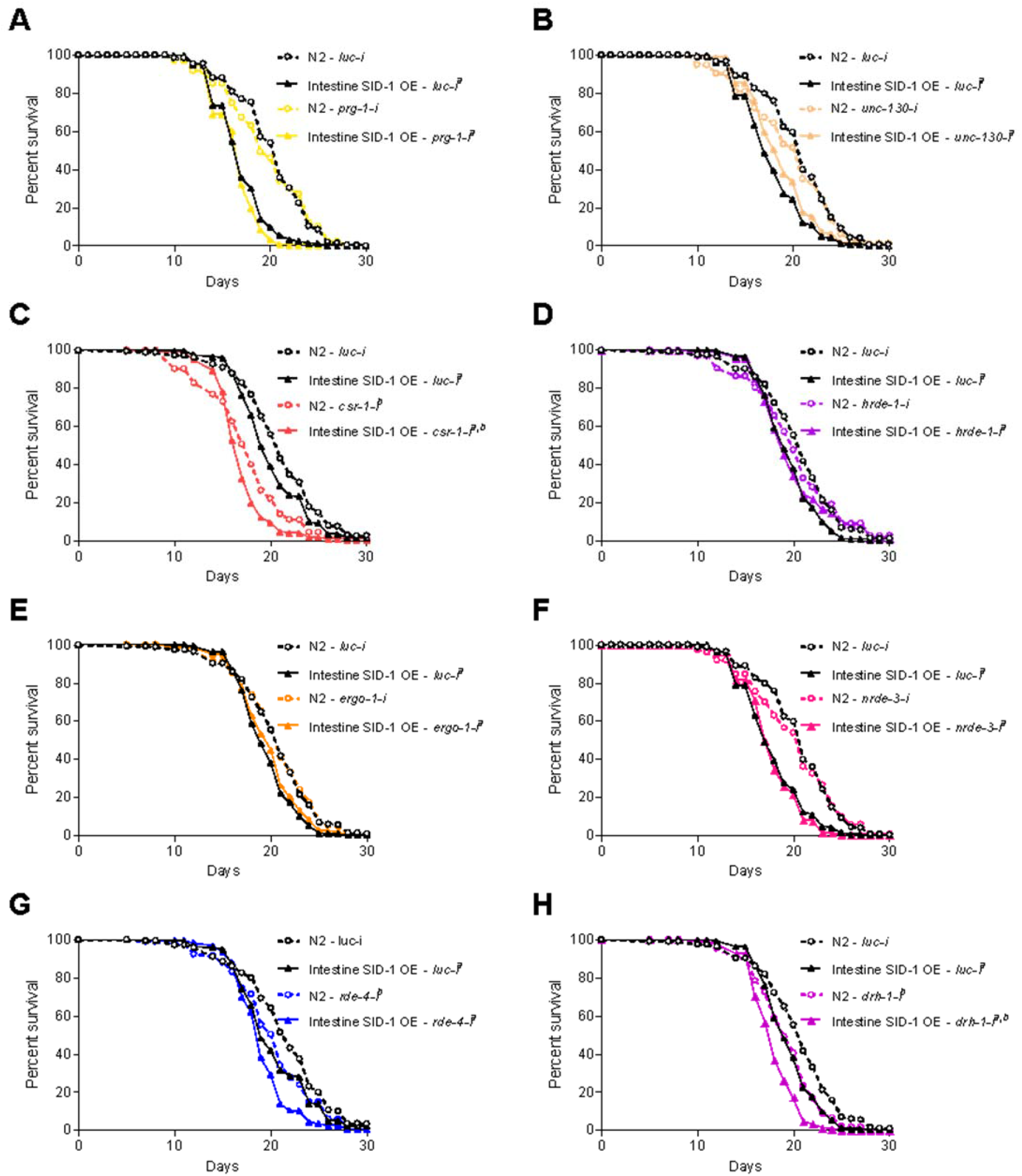
RNA pathways not involved in lifespan reduction mediated by SID-1 overexpression. Lifespan of control (N2) and intestinal *sid-1* overexpressing (*gly-19p::sid-1*) animals maintained at 20°C and fed the RNAi targeting luciferase or *prg-1* (A), *unc-130* (B), *csr-1* (C), *hrde-1* (D), *ergo-1* (E), *nrde-3* (F), *rde-4* (G), and *drh-1* (H). Lifespan was compared by the log-rank test (^a^P<0.05 in comparison to wild-type control; ^b^P<0.05 in comparison to luciferase RNAi control). Data are the average of two (D, E) and three (A-C, F-H) independent trials. See Supplementary Table 1 for lifespan statistics.

### The bacterial source influences the lifespan outcome of tissue-specific *sid-1* overexpression models

The observation that *sid-2* RNAi partially rescued the lifespan of worms overexpressing *sid-1* in the intestine (Figure 3C) indicated that exogenous bacterial RNA contributes to reduced longevity in this model. This was corroborated by the observation that the average lifespan reduction produced by intestinal *sid-1* overexpression was less evident in our RNAi screen, performed with the RNAi-compatible *E. coli* HT115(DE3), in comparison to our initial experiments performed using OP50-1. Specifically, the mean lifespan decrease of intestinal *sid-1* overexpressing worms was reduced from 24.0 ± 1.5% in the *E. coli* OP50-1 to 9.2 ± 6.2% in the *E. coli* HT115(DE3).

To directly compare the effects of the bacterial diet on the lifespan reduction induced by tissue-specific *sid-1* overexpression, we exposed worms to HT115(DE) or OP50(*xu363*). This RNAi-compatible OP50 strain shares the RNase III mutation and tetracycline resistance with HT115(DE3) (Xiao *et al*., 2015). Therefore, lifespan assays were conducted in parallel in tetracycline-supplemented plates, isolating the bacterial source as the sole variable between conditions. While the different bacteria did not affect the lifespan of wild-type worms (Figure 4), we observed that the lifespan of worms overexpressing *sid-1* in the intestine was shorter in OP50(*xu363*) than in HT115(DE) (Figure 4), showing that the bacterial source determines the magnitude of lifespan reduction produced by *sid-1* overexpression.

**Figure 4.**
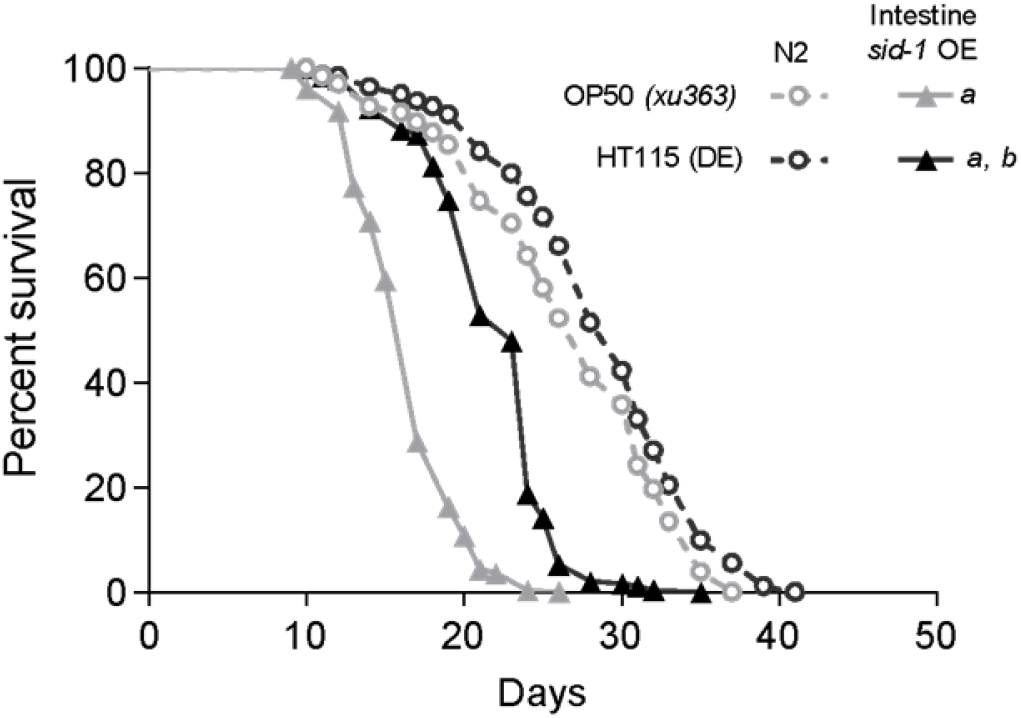
HT115 feeding attenuates the lifespan reduction of worms with intestine-specific *sid-1* overexpression. Lifespan of control and intestinal *sid-1* overexpressing (*gly-19p::sid-1*) animals fed the RNAi-competent HT115(DE) or OP50(*xu363*). Lifespan was compared by the log-rank test [^a^P<0.05 in comparison to wild-type control; ^b^P<0.05 in comparison to OP50(*xu363*)]. Data are representative of two independent trials. See Supplementary Table 1 for lifespan statistics.

### Tissue-specific SID-1 overexpression reveals a delicate balance for intercellular RNA signaling

The presence of non-coding RNAs in extracellular fluids has been long recognized (Turchinovich *et al*., 2013), but evidence of cell non-autonomous gene regulation by these circulating species has only recently been described (Thomou *et al*., 2017; Ying *et al*., 2017; Zhang *et al*., 2017; Mori *et al*., 2019; Zhou *et al*., 2019). In the present work, we observed reduced lifespan by increasing the tissue-specific expression of the RNA transporter SID-1 in *C. elegans*. The lifespan reduction produced by tissue-specific SID-1 overexpression depends on components of the systemic RNAi and miRNA pathways, implicating extracellular non-coding RNA signaling in the phenotype. In addition, the lifespan reduction produced by intestinal *sid-1* overexpression is partially dependent on *sid-2* and altered by the bacterial diet, suggesting diet-derived environmental RNAs are potential regulators of longevity in these models.

SID-1 canonically functions as a transmembrane dsRNA-operated channel that allows the selective uptake of double-stranded structured RNAs (including single-stranded hairpins, such as pre-miRNAs) (Winston, Molodowitch and Hunter, 2002; Shih *et al*., 2009; Shih and Hunter, 2011). In *C. elegans*, low expression levels of *sid-1* in neurons make these cells refractory to systemic RNAi (Calixto *et al*., 2010). Accordingly, overexpression of *sid-1* in neurons restores their sensitivity to environmental RNAi while reducing RNAi efficiency in other tissues (Calixto *et al*., 2010), suggesting that the intercellular function of RNAs follows a stoichiometric balance greatly influenced by the expression level of SID-1 across tissues. Indeed, we observed reduced lifespan in *C. elegans* models of dysregulated intercellular RNA signaling by overexpression of SID-1 in specific tissues. The lifespan reduction of tissue-specific *sid-1* overexpressing worms is attenuated by silencing components of the systemic RNAi pathway, confirming the requirement of intertissue RNA transport. Based on these results, we propose that enhanced uptake of extracellular RNAs induced by SID-1 overexpression affects the stoichiometric balance between intracellular and extracellular RNAs, thereby dysregulating gene expression. We termed this model <u>In</u>tracellular/<u>Ex</u>tracellular <u>S</u>ystemic RNA imbalance (InExS) (Figure 5).

**Figure 5.**
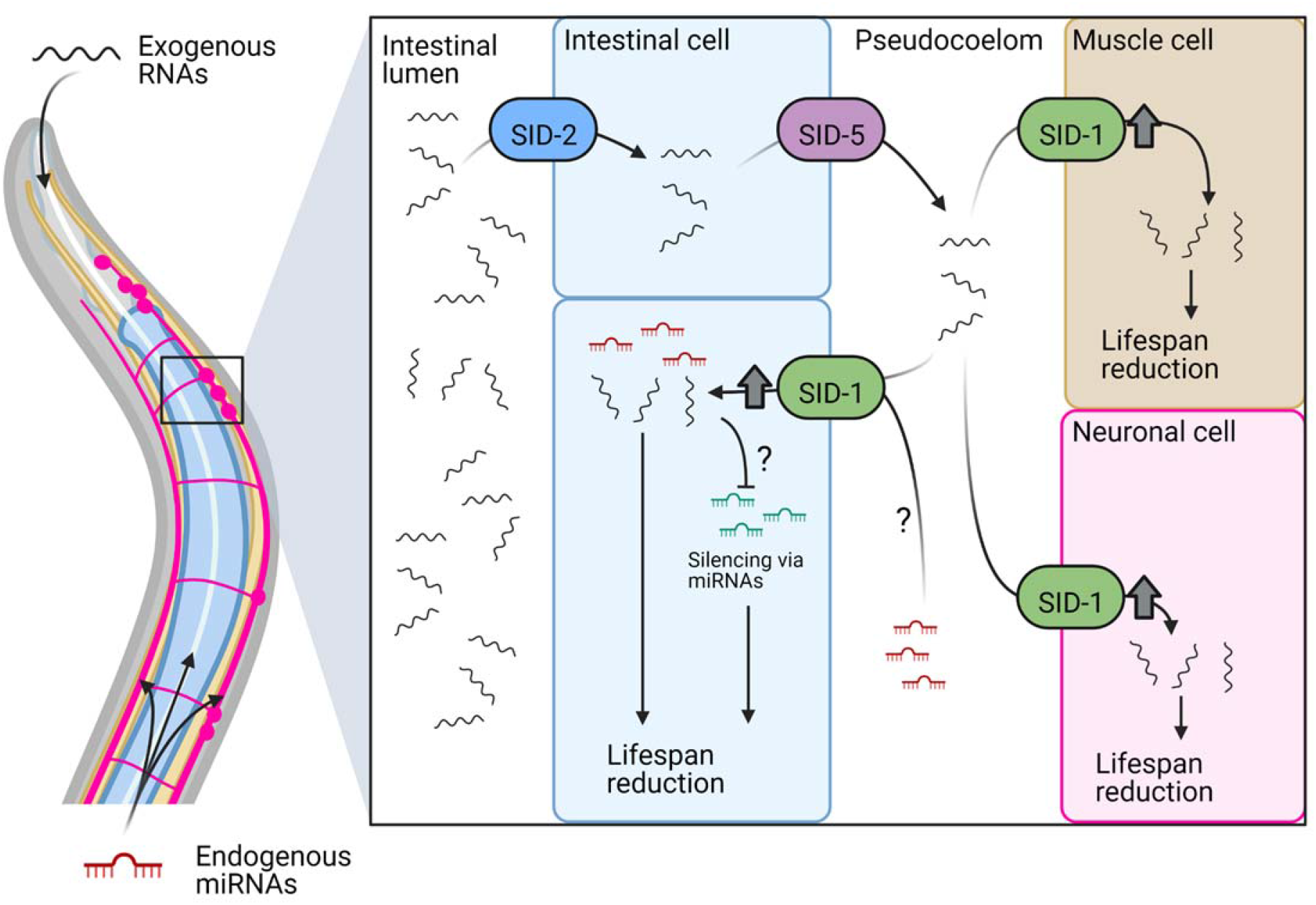
<u>In</u>tracellular/<u>Ex</u>tracellular <u>S</u>ystemic RNA imbalance (InExS) caused by tissue-specific SID-1 overexpression. SID-1 overexpression in the intestine, muscle or neurons causes lifespan reduction. In the intestinal SID-1 overexpression model, this reduction is partially mediated by environmental RNAs internalized via SID-2 and exported to the pseudocoelom by SID-5. Increased SID-1-mediated uptake of RNAs causes an imbalance between intracellular and extracellular RNAs impairing the cell-autonomous function of non-coding RNAs such as miRNAs. Created with BioRender.com.

Competition between RNA species for rate-limiting shared enzymes of different small RNA pathways has been previously reported (Simmer *et al*., 2002; Timmons *et al*., 2003; Calixto *et al*., 2010; Jimmy and Hunter, 2011; Jose *et al*., 2012) and disruption of this balance is physiologically relevant. For example, embryonic lethality induced by loss of the miR-35-41 family (Massirer *et al*., 2012; De-Souza *et al*., 2019) – a family of miRNAs that are highly expressed during development - is reversed by environmental RNAi even in the absence an endogenous target (e.g., feeding luciferase RNAi) (De-Souza *et al*., 2019), demonstrating that the miRNA and systemic RNAi pathways are tightly intertwined. In our experiments, we observed that intestinal *sid-1* overexpression results in lifespan reduction which is not additive to the reduction promoted by *dcr-1* or *pash-1* RNAi, indicating that impaired miRNA function is involved in the mechanism of lifespan reduction by InExS.

### Bacterial diet contributes to lifespan reduction induced by InExS

We also found that the lifespan reduction caused by intestinal *sid-1* overexpression is attenuated by *sid-2* RNAi (Figure 3). SID-2 is essential for the uptake of environmental RNAs via the intestine (Winston *et al*., 2007), therefore implicating bacterial RNAs in the lifespan reduction resulting from InExS. Consistently, we observed that feeding two distinct RNAi-compatible *E. coli* strains [OP50(*xu363*) and HT115(DE)] differentially affected the lifespan of *sid-1* overexpressing worms, but not wild-type controls (Figure 4). Worms are constantly surveilling dietary RNA composition and responding to it (Kaletsky *et al*., 2020; Moore *et al*., 2021). The *E. coli* non-coding RNA DsrA was shown to reduce *C. elegans* lifespan in *sid-1-* and *sid-2-*independent manners (Liu *et al*., 2012) and without accumulation of primary and secondary siRNAs (Akay, Sarkies and Miska, 2015), suggesting no activation of the canonical RNAi pathway. In contrast, other reports have shown that exposure to pathogenic *Pseudomonas aeruginosa* (PA14) elicits an avoidance response that is mediated by the RNAi pathway (Moore, Kaletsky and Murphy, 2019; Kaletsky *et al*., 2020). Interestingly, exposure to a single small non-coding RNA from PA14 is both necessary and sufficient to induce avoidance behavior, and it does so in *sid-2-* and *dcr-1-* dependent manners (Kaletsky *et al*., 2020). On the other hand, *sid-1* is required for the transgenerational inheritance of this phenotype. These mechanisms of RNA surveillance have been proposed to act as a way of mapping the worm bacterial environment in a signal transmittable throughout generations (Braukmann, Jordan and Miska, 2017; Manterola, Palominos and Calixto, 2021; Chen and Rechavi, 2022). Indeed, our data support the notion that RNAs from different bacterial sources can contribute differently to InExS, thus reducing worm lifespan when there is tissue-specific overexpression of SID-1.

In the present work, we interfered with the systemic dsRNA transport machinery in *C. elegans* by manipulating the expression of SID-1 in different tissues. Dysregulation of RNA transport manifested as a reduction in lifespan, which is mechanistically linked to the ingestion of bacterial RNAs and associated with impaired miRNA function. These observations led us to propose that intertissue RNA signaling is tightly regulated in *C. elegans* and the imbalance between intracellular and extracellular RNAs promoted by tissue-specific *sid-1* overexpression and the uptake of environmental RNA reduces worm lifespan. Further studies are required to assess the specific RNAs eliciting these phenotypes and how translatable these mechanisms are to humans, although tissue-specific Dicer knockout in mice and downregulation of the circulating exosomal miRNA pool in mice and humans have already been associated with metabolic dysfunction and age-related features (Reis *et al*., 2016; Thomou *et al*., 2017), raising the intriguing possibility that changes in RNA transport can also accelerate aging in mammals.

## ACKNOWLEDGEMENTS

We thank Elzira E. Saviani for technical support and the *C. elegans* Genetics Center (University of Minnesota) - funded by the NIH Office of Research Infrastructure Programs (P40 OD010440) - for providing *C. elegans* strains. We also thank Adam Antebi (Max Planck Institute for Biology of Aging) and Julio Ferreira (University of São Paulo) for RNAi clones. This work was funded by the Conselho Nacional de Desenvolvimento Científico e Tecnológico (CNPq) – grant number 310287/2018-9 (M.A.M.), Coordenação de Aperfeiçoamento de Pessoal de Nível Superior - Brasil (CAPES) - Finance Code 001, 88881.143924/2017-01 (M.A.M.), and 88887.489628/2020-00 (G.T.S), and PROEX0230081 (H.C.), and Fundação de Amparo à Pesquisa do Estado de São Paulo (FAPESP) - grant numbers 2019/25958-9 (M.A.M.), 2017/01184-9 (M.A.M.), 2014/10814-8 (E.A.S.), 2019/01587-1 (C.A.V.J), 2019/04726-2 (T.L.K.), 2017/22057-5 (W.G.S.), 2016/05560-2 (J.R.), 2015/04264-8 (H.C), and 2017/01339-2 (H.C.).

## CONFLICT OF INTEREST

The authors declare no conflict of interest.

## AUTHOR CONTRIBUTION

**Henrique Camara:** Conceptualization, Formal analysis, Investigation, Writing - Original Draft, Visualization, Project administration; **Carlos A. Vergani-Junior:** Investigation, Validation, Writing - Review & Editing; **Silas Pinto:** Conceptualization, Investigation, Validation; **Thiago L. Knittel:** Investigation; **Willian G. Salgueiro:** Investigation, Validation; **Guilherme Tonon-da-Silva:** Investigation; **Juliana Ramirez:** Investigation; **Evandro A. De-Souza:** Conceptualization, Writing - Review & Editing; **Marcelo A. Mori:** Conceptualization, Writing - Review & Editing, Project administration, Funding acquisition.

## METHODS

### C. elegans

Nematode *C. elegans* were obtained from the Caenorhabditis Genetics Center (CGC). Worms were cultivated in Petri dishes containing semi-solid Nematode Growth Media (NGM) supplemented with 1 mM CaCl_2_, 1 mM MgSO_4_, 25 mM KPO_4_ and 5 μg/ml cholesterol. To avoid contamination, the NGM medium was supplemented with 100 μg/ml of streptomycin for the maintenance plates seeded with OP50-1. RNAi plates seeded with HT115(DE) were supplemented with 1 mM CaCl_2_, 1 mM MgSO_4_, 25 mM KPO_4_, 8 μg/ml cholesterol and12.5 μg/ml of tetracycline and 100 μg/ml of ampicillin as the antibiotics. RNAi plates were also supplemented with 1 mM IPTG to induce double-stranded RNA expression. For the assay comparing HT115(DE) and OP50(*xu363*), plates were supplemented with 1 mM CaCl_2_, 1 mM MgSO_4_, 25 mM KPO_4_, 8 μg/ml cholesterol and 12.5 μg/ml of tetracycline. Bacteria were grown for 16 hours, during which they reached an optical density of approximately 1. Then, the culture was concentrated 10 times and inoculated onto plates containing NGM, forming a layer under which the worms were cultivated. For most experiments, worms were kept at 20°C, except for the heat stress protocol, where maintenance was carried out at 28°C from day 0 of adulthood. Table 1 summarizes the lineages used in the present work.

**Table 1.**
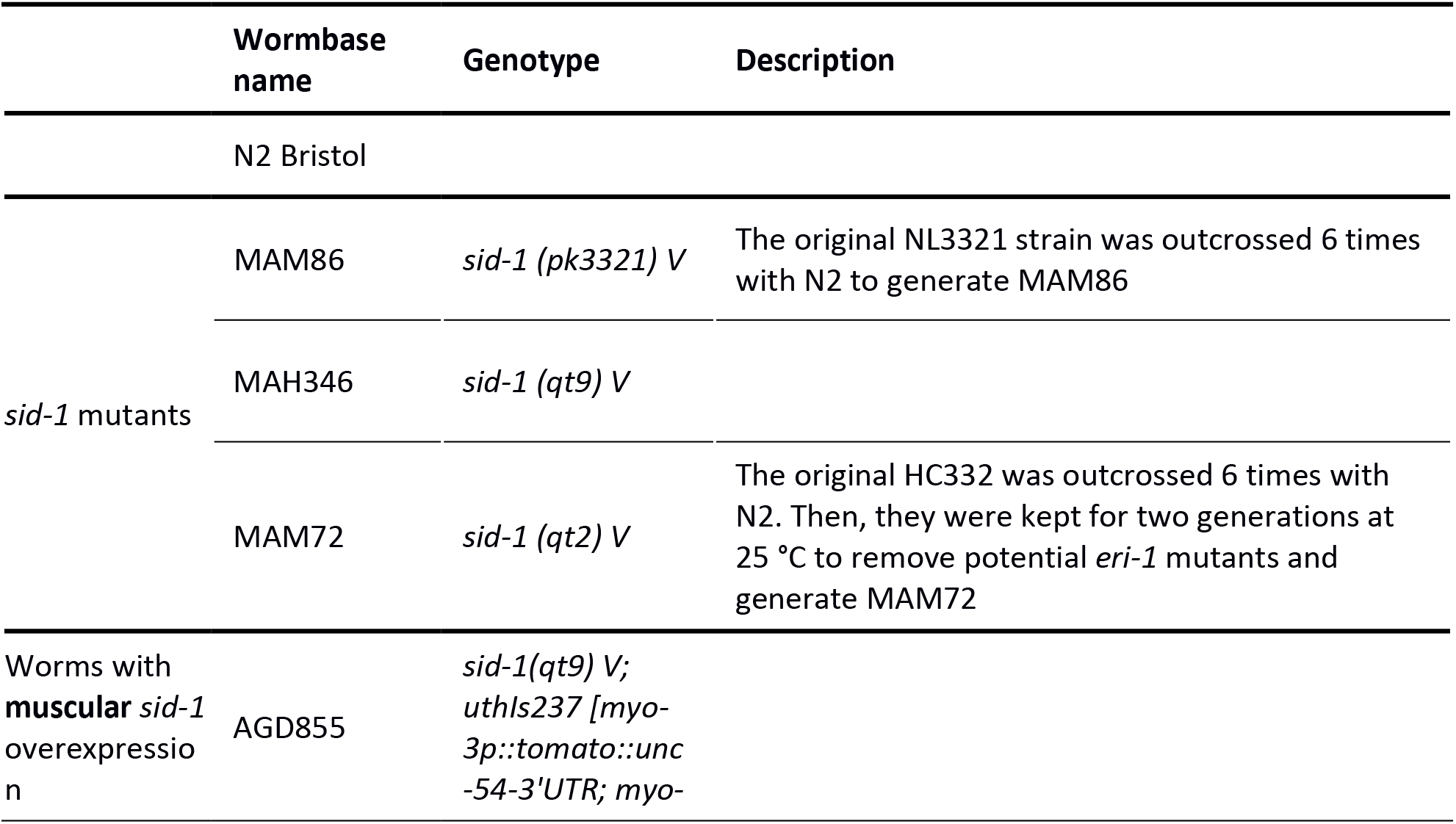
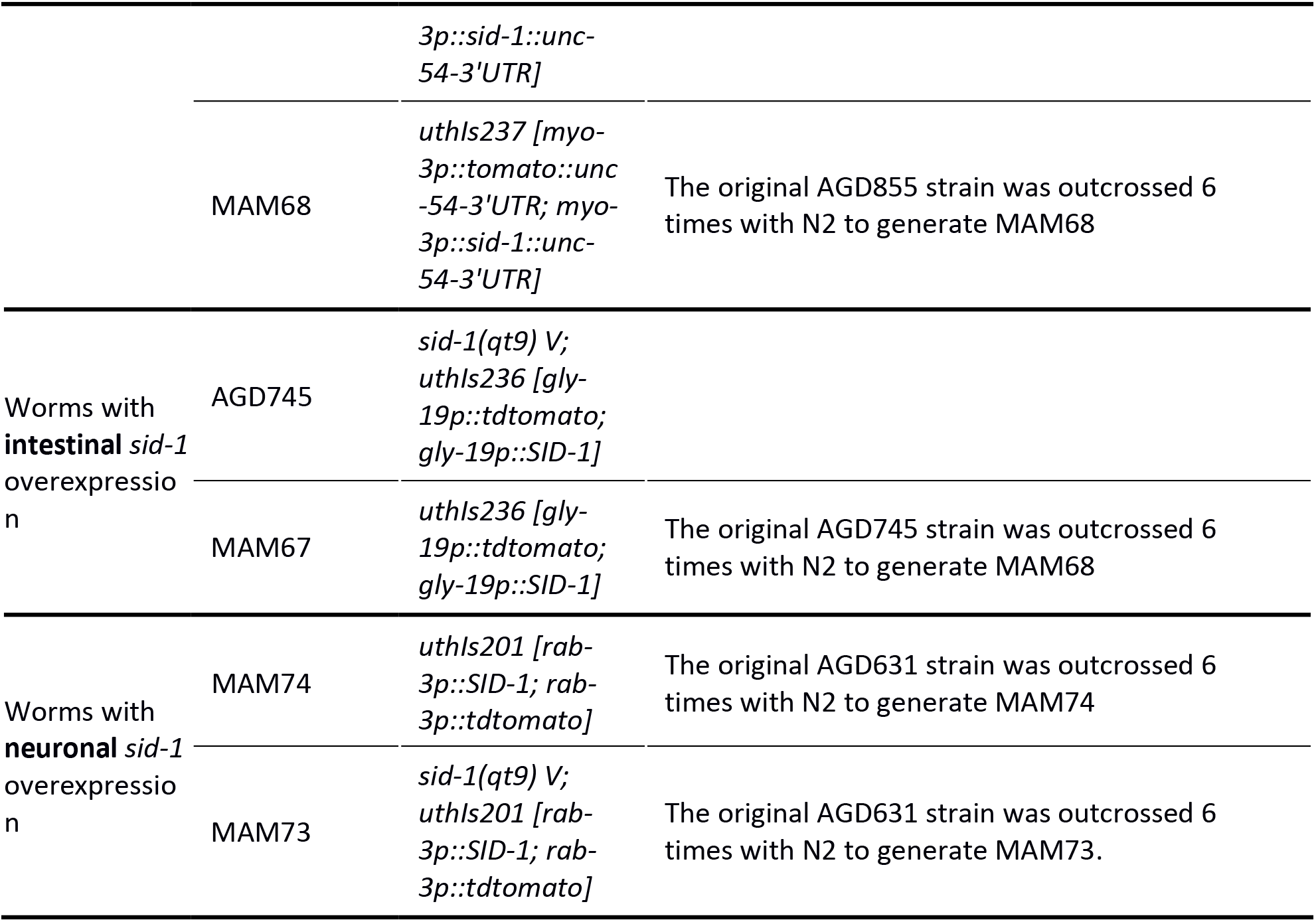
*C. elegans* strains used in the present work.

### RNAi clones

RNAi clones were from the Vidal RNAi library, kindly provided by Dr. Adam Antebi and Dr. Julio Ferreira, with exception of *sid-1* RNAi. The construction of RNAi for *sid-1* was similar to the protocol described before (Ferraz *et al*., 2016). In summary, part of an exon of the *sid-1* gene was amplified by PCR from wild-type genomic DNA of worms using the primer 5’-AACCATGGATGGTCCGTCAGTGGTTTGT-3’ and the primer 5’-TACTCGAGAGCAGTGACAGAGCTCCACA-3’ (Exxtend). The PCR conditions were: 95°C for 5 min; 40 cycles of 95°C for 10s, 60°C for 30s and 72°C for 30s; 72°C for 5 min. The PCR product was loaded on a 1% agarose gel followed by electrophoresis to purify the expected product. The product was extracted from the gel using the QIAquick Gel Extraction Kit (QIAGEN) according to the manufacturer’s recommendations. Approximately 1 μg of the purified product was used as well as the target vector L4440 for digestion with restriction enzymes NcoI and XhoI for 3h at 37°C in NEB 2 buffer following the manufacturer’s recommendations (New England Biotechnology). Ligation of the insert to the vector was performed using the Quick Ligation kit (New England Biolabs). The final vector was transformed into *E. coli* TOP10 (One Shot^®^ iTOP10 Chemically Competent E. coli) and recombinant colonies were selected for ampicillin resistance. Positive recombinants were identified by colony PCR using the same primers used for *sid-1* exon amplification. The plasmid was purified using the QIAprep Spin Miniprep Kit (QIAGEN) and validated by sequencing. After sequence confirmation, the plasmid was used to transform *E. coli* HT115(DE3). Recombinants were selected for resistance to ampicillin and tetracycline, expanded, and used for RNAi experiments in worms. *Sid-1* knock-down efficiency was confirmed by qPCR in animals fed *sid-1* or *luc-i* RNAi from hatching to D0 of adulthood (Supplementary figure 3).

**Supplementary Figure 3.**
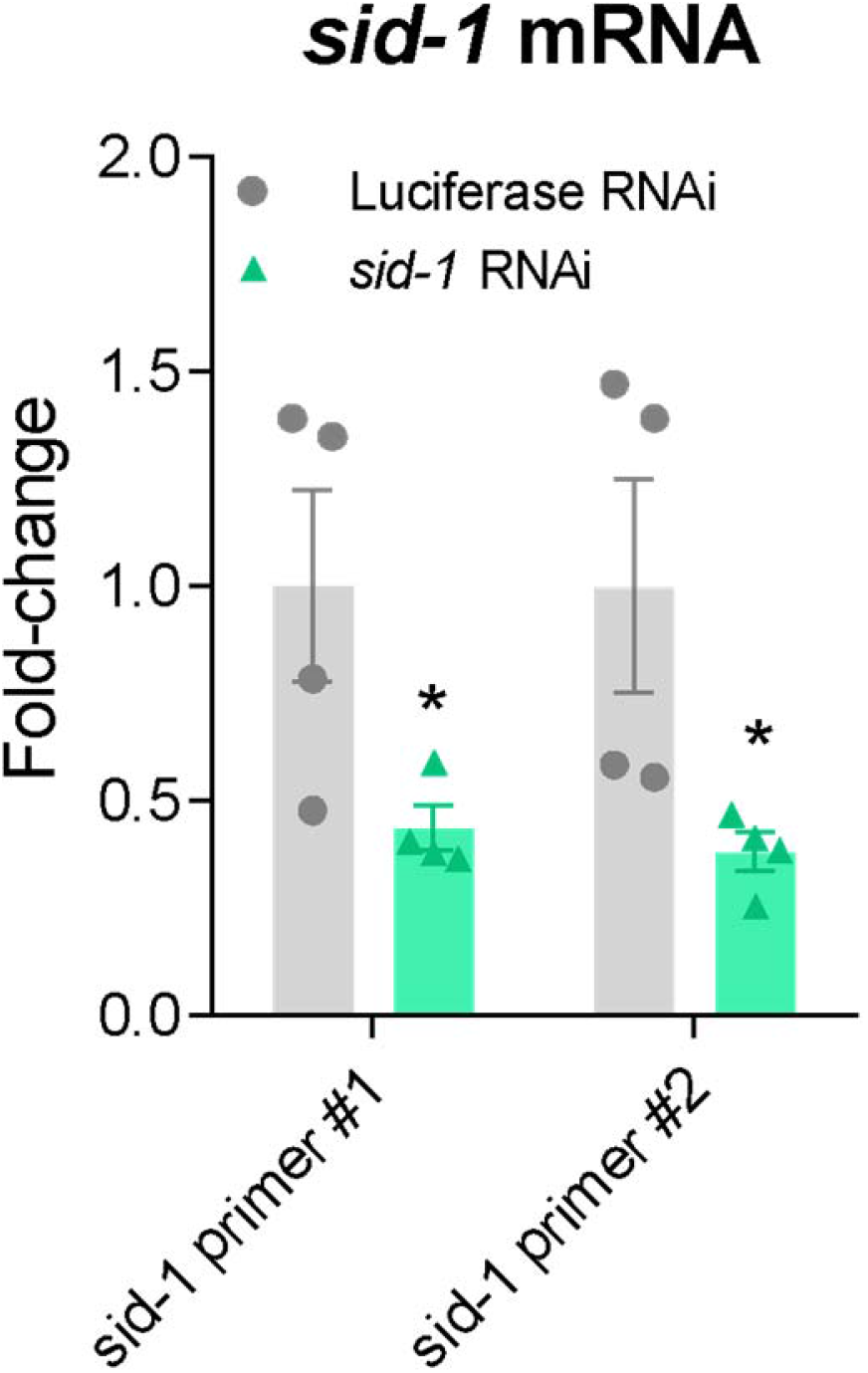
Validation of *sid-1* RNAi efficiency. Expression of *sid-1* on D0 adult control (N2) fed with bacteria expressing RNAi against luciferase or *sid-1* from hatching (*P<0.05, n=4, Multiple t-test).

### Lifespan

To quantify the lifespan of *C. elegans*, synchronized populations were obtained by bleaching gravid hermaphrodites and plating the eggs in NGM plates with food. For the RNAi lifespan assays, worms hatched in plates seeded with the corresponding RNAi to induce gene knock-down during their whole life. Approximately 150 synchronized L4 worms were transferred to plates supplemented with 0.5 μg/ml of 5-fluor-2-deoxyuridine - FUdR (Sigma), an inhibitor of mitosis, to prevent offspring development and overpopulation (Olsen, Vantipalli and Lithgow, 2006). From then on, each strain was monitored every other day for identification and removal of dead worms. The worms were classified as dead due to the lack of motor response to touch with the platinum pick at their tail and head. Plates were monitored until the identification of the last dead worm. For the thermic stress experiment, worms were transferred to 28°C incubators at day 0 of adulthood. Otherwise, experiments were carried out at 20°C.

### Oxidative stress

The acute oxidative stress protocol was performed by adding 200 μl of sodium arsenite at 0 mM, 7.5 mM, 10 mM or 20 mM diluted in M9 medium (3 g KH_2_PO_4_, 6 g Na_2_HPO_4_, 5 g NaCl, 1 ml 1 M MgSO_4_, H_2_O to 1 liter, sterilized by autoclaving) in the wells of a 96-well plate. For each concentration of sodium arsenite, worms from different strains were distributed at a density of 5-15 worms/well on day 0 of adult life and kept at 20°C for 6-8 hours. After this period, the number of surviving worms was evaluated by inspecting signs of motor activity with the aid of a light stereoscope.

### Egg Laying

As an estimation of fertility, total egg laying was performed as in Ferraz et al. 2016 (Ferraz *et al*., 2016). In summary, eight L4 larval stage hermaphrodites were individualized in one well of a 24-well plate containing NGM seeded with 10 μl of OP50-1. Each day, the hermaphrodites were transferred to a new well and the laid eggs were counted. Progeny viability in each well was evaluated by quantifying larvae hatched 48 hours after hermaphrodite removal from that well. Egg laying was scored for 10 days to cover the whole reproductive period of the hermaphrodites and the total amount of eggs was calculated and compared between the lines.

### Developmental time

To determine the development time, twenty young hermaphrodites were transferred to plates containing NGM with OP50-1 at 20°C for 2 hours for laying eggs. Then, the hermaphrodites were removed and the plates were kept at 20°C for 60 hours. From hour 60 onwards, the plates were monitored every hour for the identification and removal of adult worms identified by the presence of eggs in the gonads. The number of adult worms at each observation point was used to generate the development curve for each strain (Ferraz *et al*., 2016).

### qPCR

For RNA extraction, worms were collected by washing the plate with 1 ml of MiliQ H_2_O and transferred to a conical tube. After decanting, the supernatant was removed and 500 μl of MiliQ water was added. This process was repeated three times to remove transferred bacteria during plate washing. 500 μl of Trizol was added to the worm pellet. To lyse the worms, magnetic beads were added to the solution and the lysis process was carried out in a magnetic stirrer. The RNA extraction was performed by adding chloroform 1/5 (v/v) to the Trizol-containing tube followed by vigorous agitation. The samples were kept at room temperature for 10 minutes and then centrifuged at 12,000 x g for 15 min at 4°C. The upper phase of the solution was transferred to a new tube. Isopropyl alcohol 1/1 (v/v), 15 μg/ml of GlycoBlue (ThermoFisher) and 0.3 M of sodium acetate were added to the solution and then kept at −20°C overnight for RNA precipitation. After centrifugation of 12,000 x g, 10 min, 4°C the supernatant was removed. The pellet was washed with 75% ethanol, vortexed briefly and then samples were centrifuged at 7,600 x g, 5 min, 4°C. The supernatant was discarded and the pellet containing the RNA dried at room temperature. After drying, the pellet was resuspended in sterile water. The reverse transcription reaction for cDNA synthesis was performed with the High-Capacity cDNA Reverse Transcription kit (Applied Biosystems) according to the manufacturer’s protocol. For gene expression quantification, realtime PCR was performed in the CFX384 thermocycler (Bio-Rad) with the following settings: 95°C-10 min, and 40 cycles of 95°C −15s, 60°C −30s, 72°C −30s. For normalization of gene expression, the constitutive *his-10* gene was used as endogenous control. The primers used for gene expression evaluation were *his-10*_Forward – GCAATTCGTCGTCCGC, *his-10_* Reverse – GACTCCACGGGTTTCCT, *sid-1_* Forward_#1 – CGGCGAATGAATCCATCTAT, *sid-1_* Reverse_#2 – AAATGTCGGCTTTTTGGTG, *sid-1_* Forward_#2 – CAGACCCTCATTTCCATCAAA and *sid-1_* Reverse_#2 – CAAATGAGTCACATGAAGCCTAT.

### Statistical Analysis

Comparison of survival and development curves was performed using the Log-rank test (Mantel-Cox). To compare the mean of quantitative variables between the two groups, Student’s t-test was used. To compare the mean of quantitative variables of more than two groups, one-way or two-way ANOVA test was used when appropriate, and multiple comparison tests were employed as indicated in the figure legends. The level of significance adopted for rejecting the null hypothesis was set at 5%.

## SUPPLEMENTARY FILES

**Supplementary Table 2.** List of genes from RNAi pathways silenced in the lifespan screen.

| Gene | Function |
| --- | --- |
| *csr-1* | 22G-RNA argonaute |
| *ergo-1* | 26G-RNA argonaute |
| *unc-130* | 21U-RNA regulator |
| *dcr-1* | microRNA e siRNA processing |
| *pash-1* | microRNA biogenesis |
| *hrde-1* | Nuclear endo-siRNA |
| *drh-1* | Antiviral RNAi helicase |
| *nrde-3* | Nuclear endo-siRNA |
| *prg-1* | piRNA argonaute |
| *rde-4* | siRNA argonaute |
| *sid-1* | dsRNA importer channel |
| *sid-5* | Cellular secretion of RNAs |
| *sid-2* | Uptake of environmental RNAs |

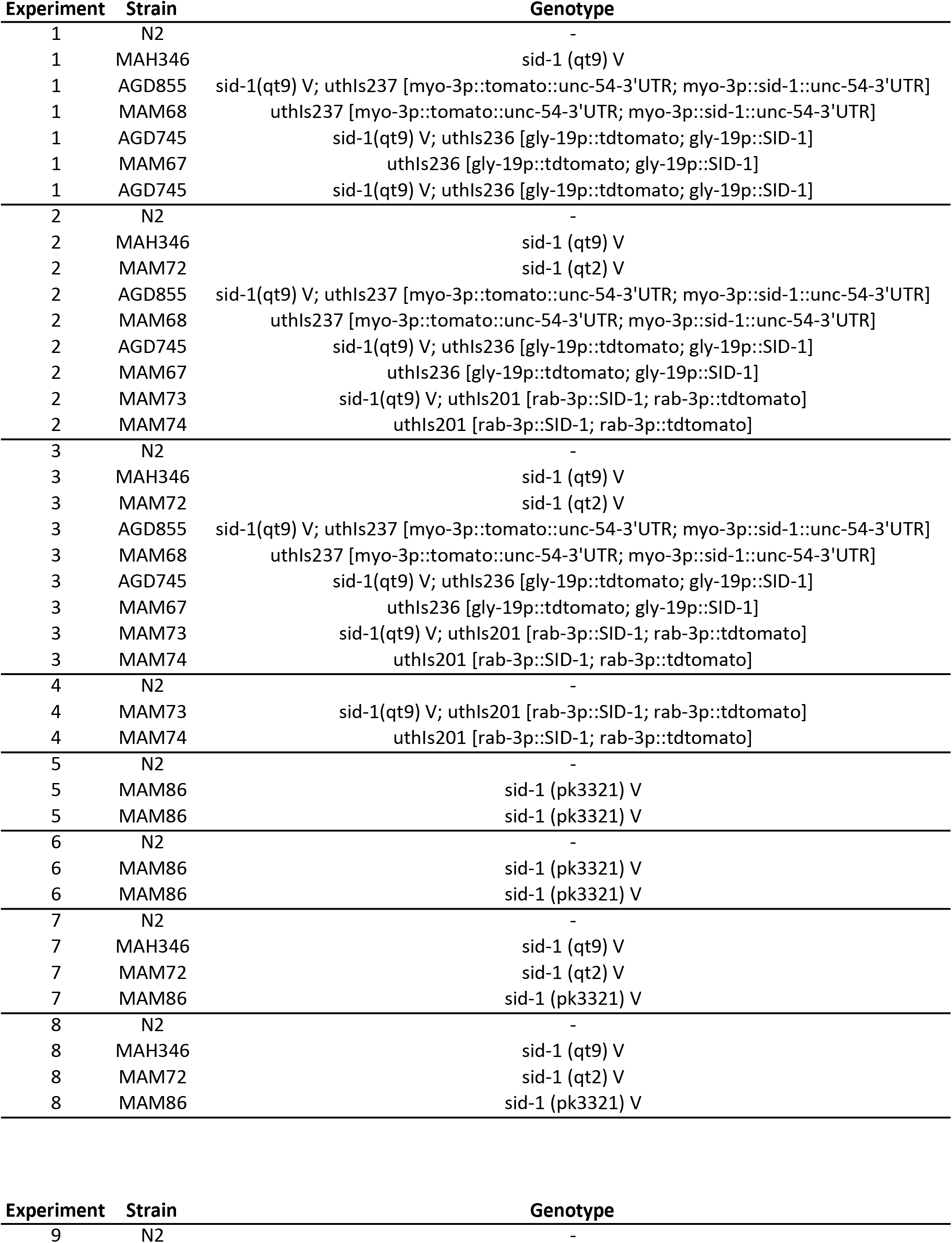
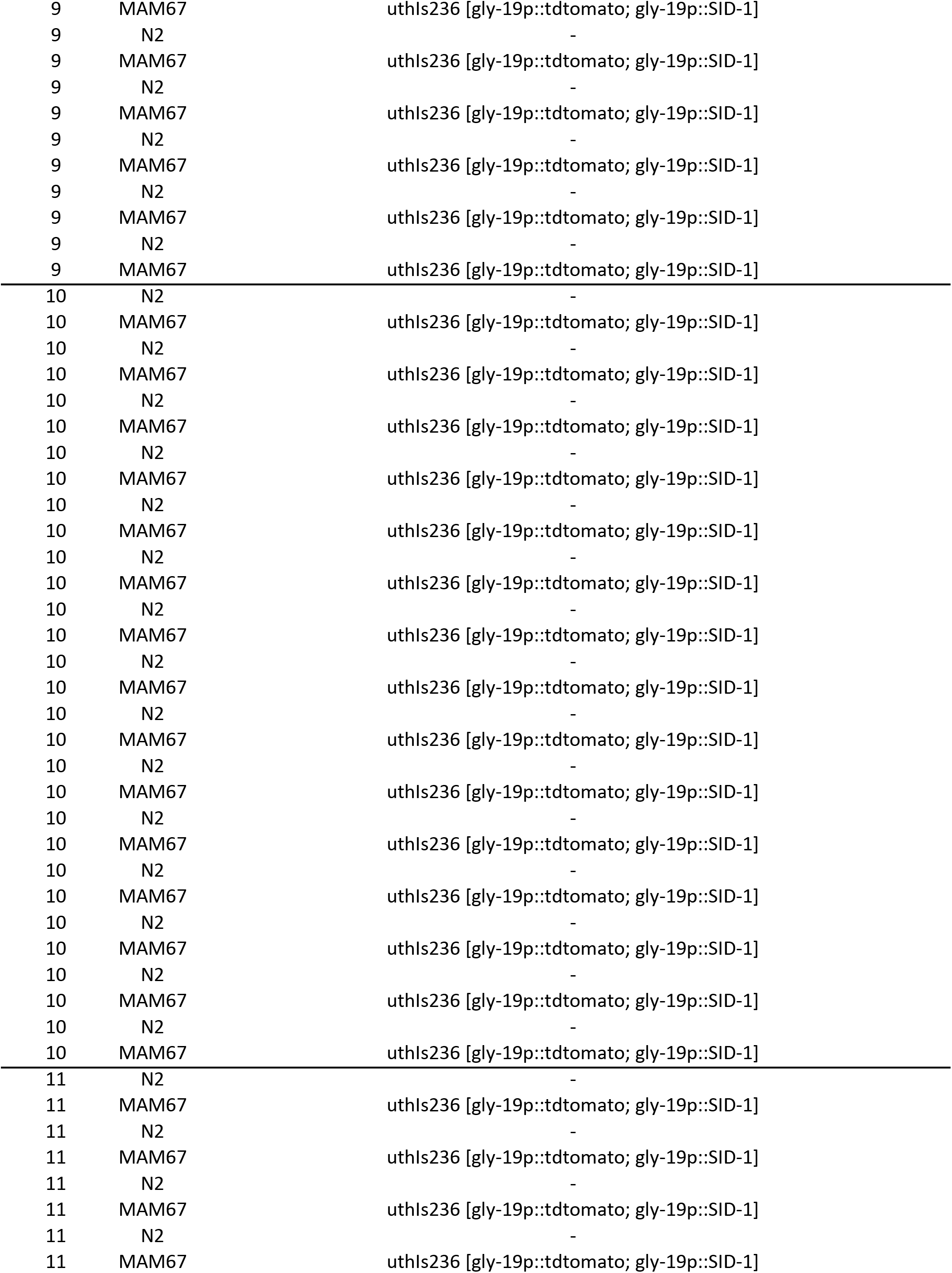
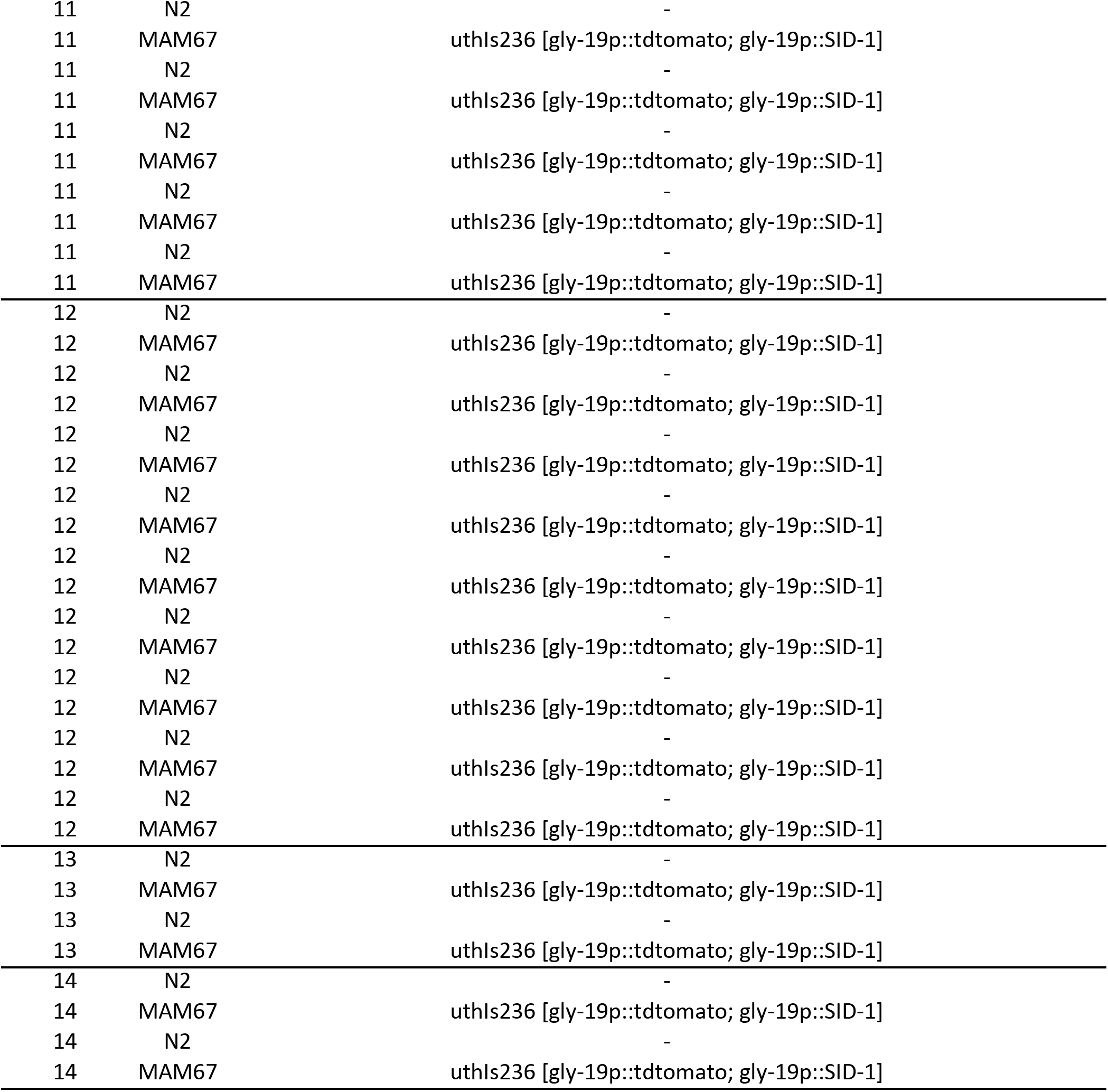
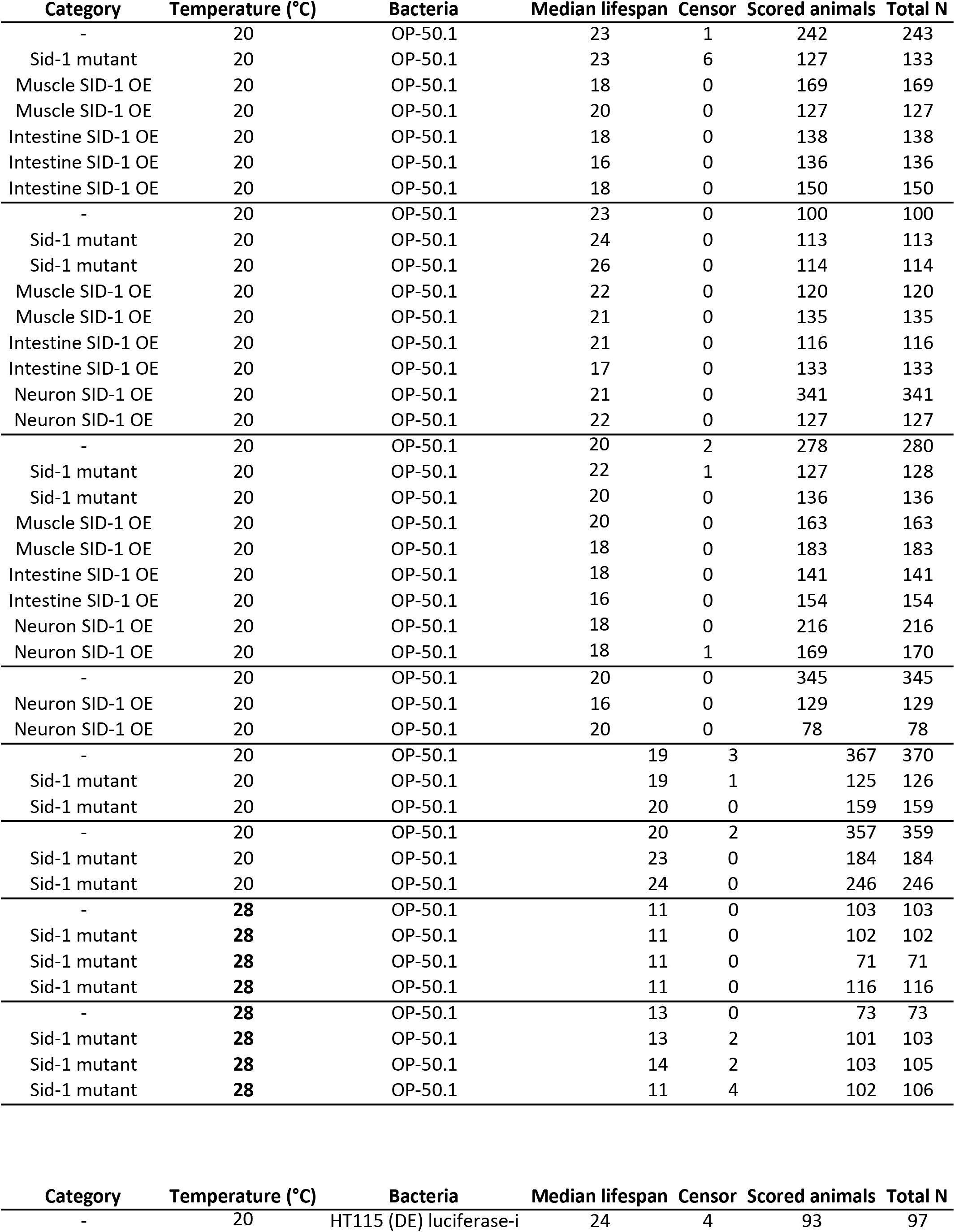
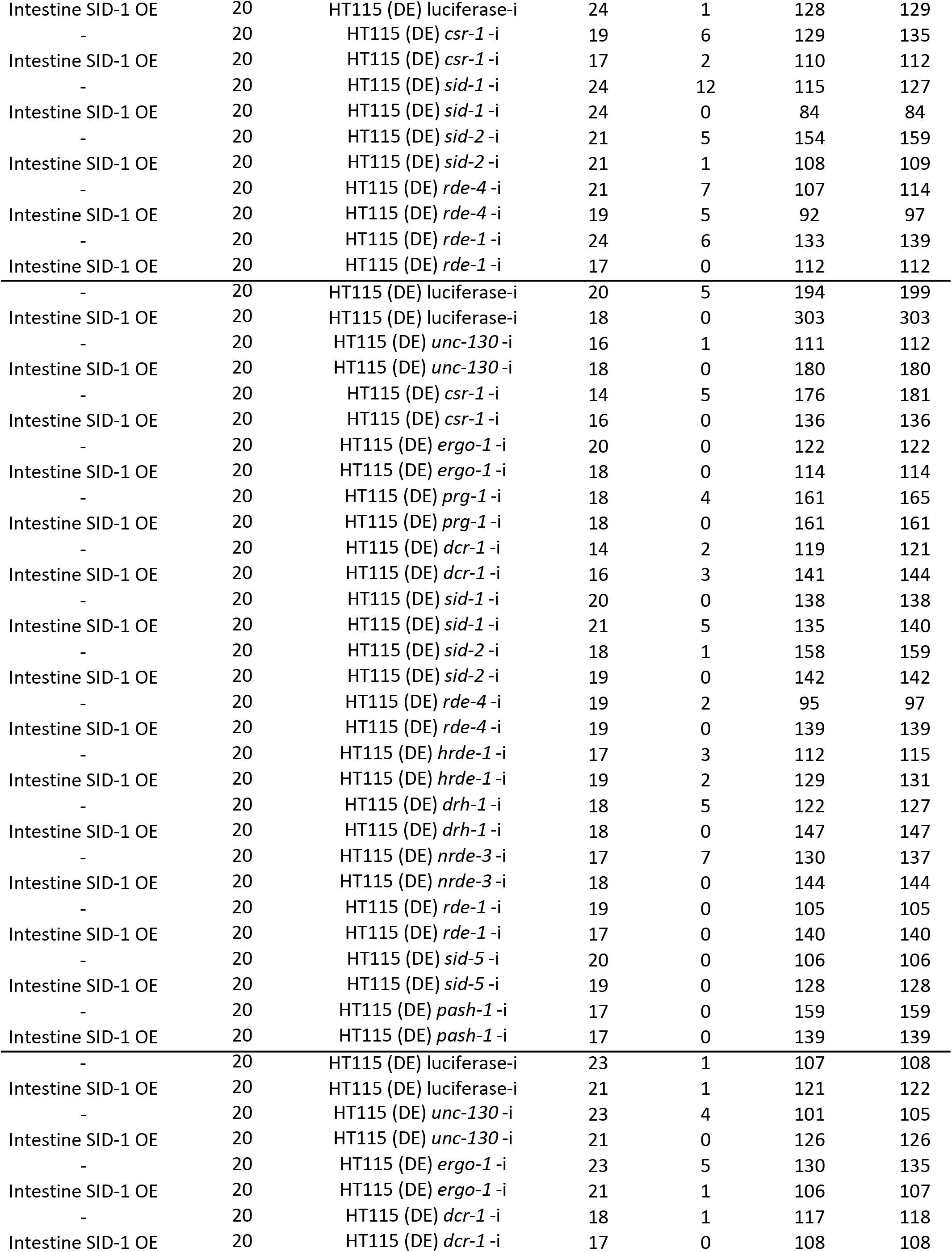
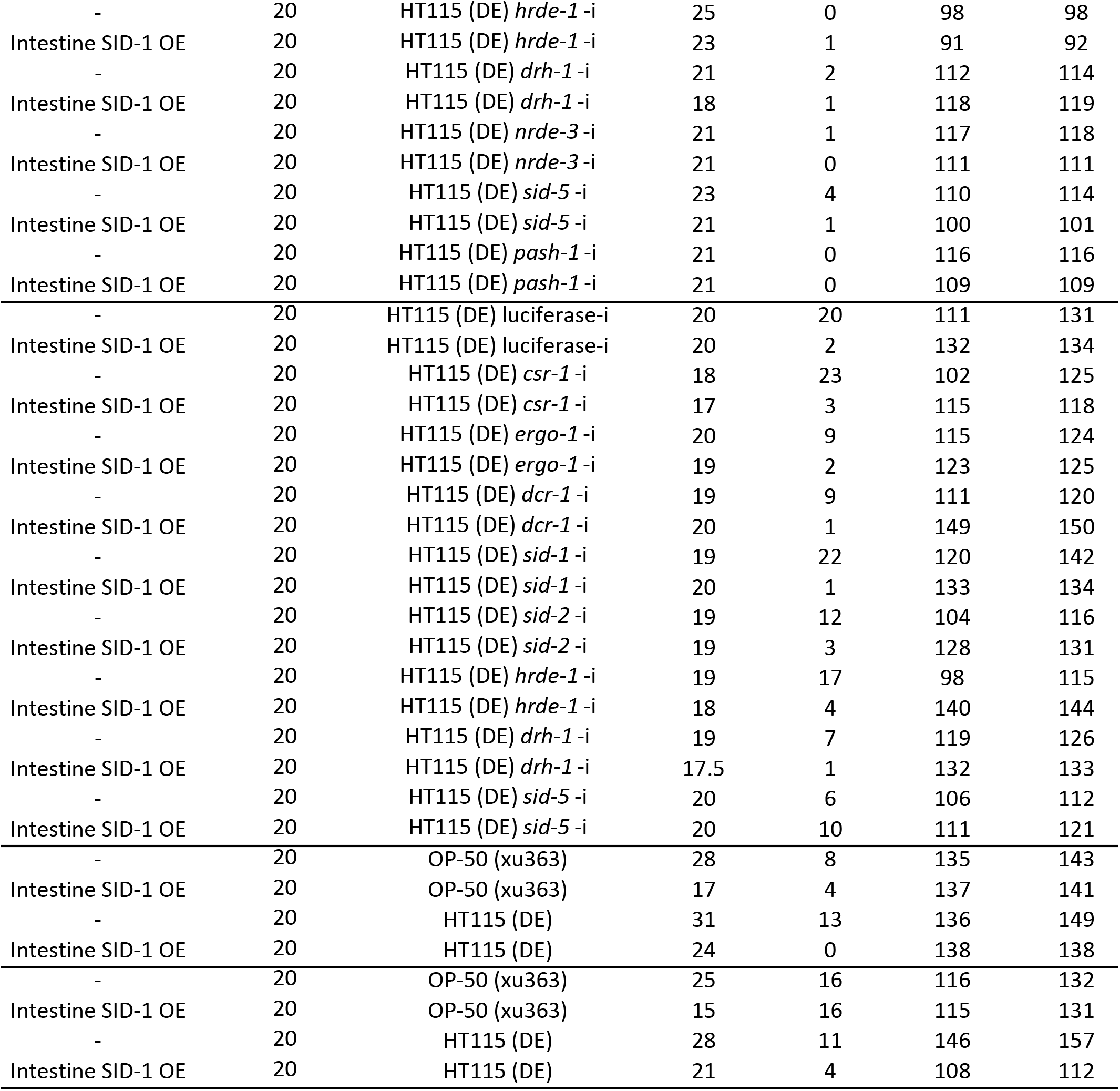
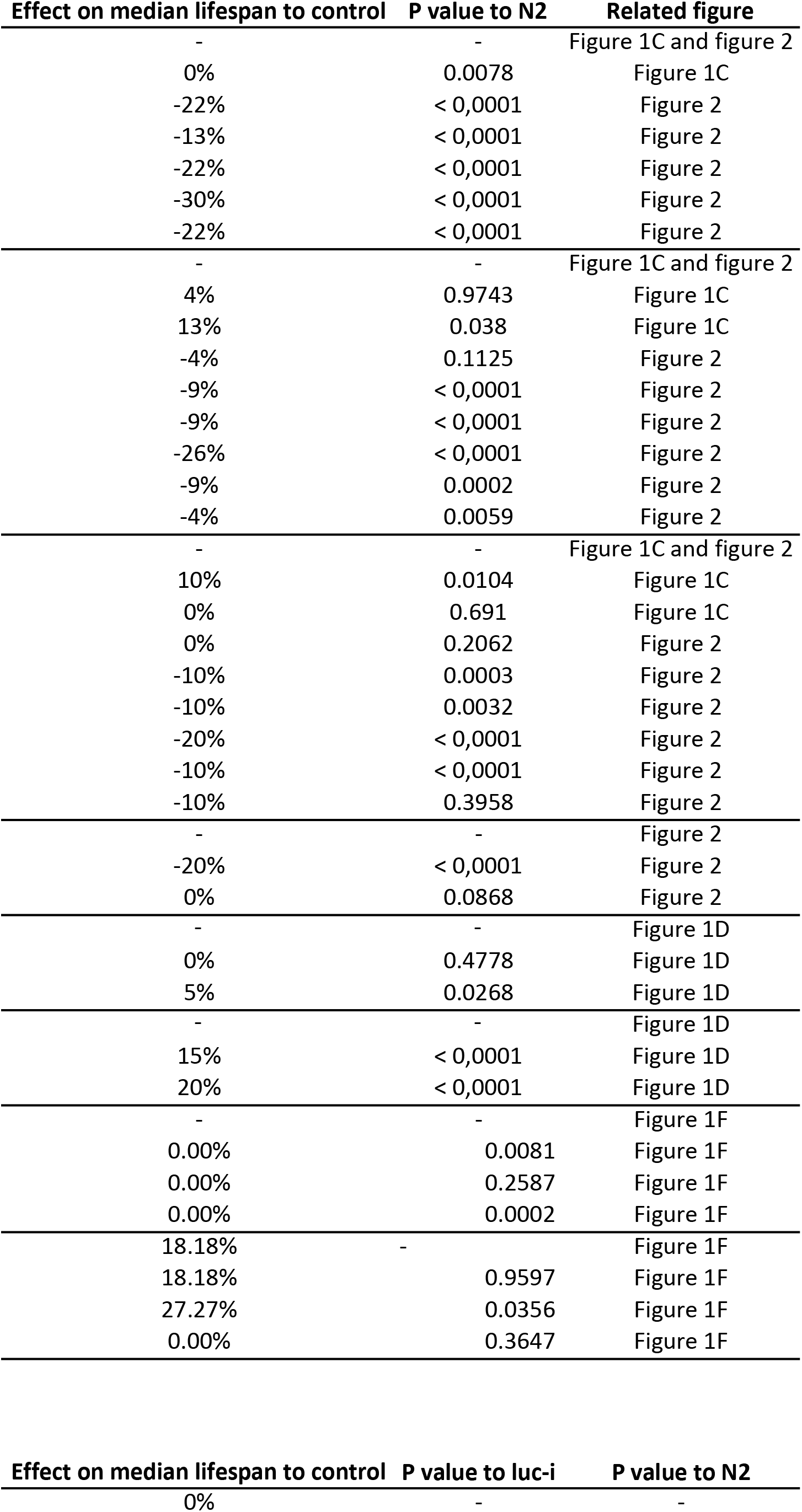
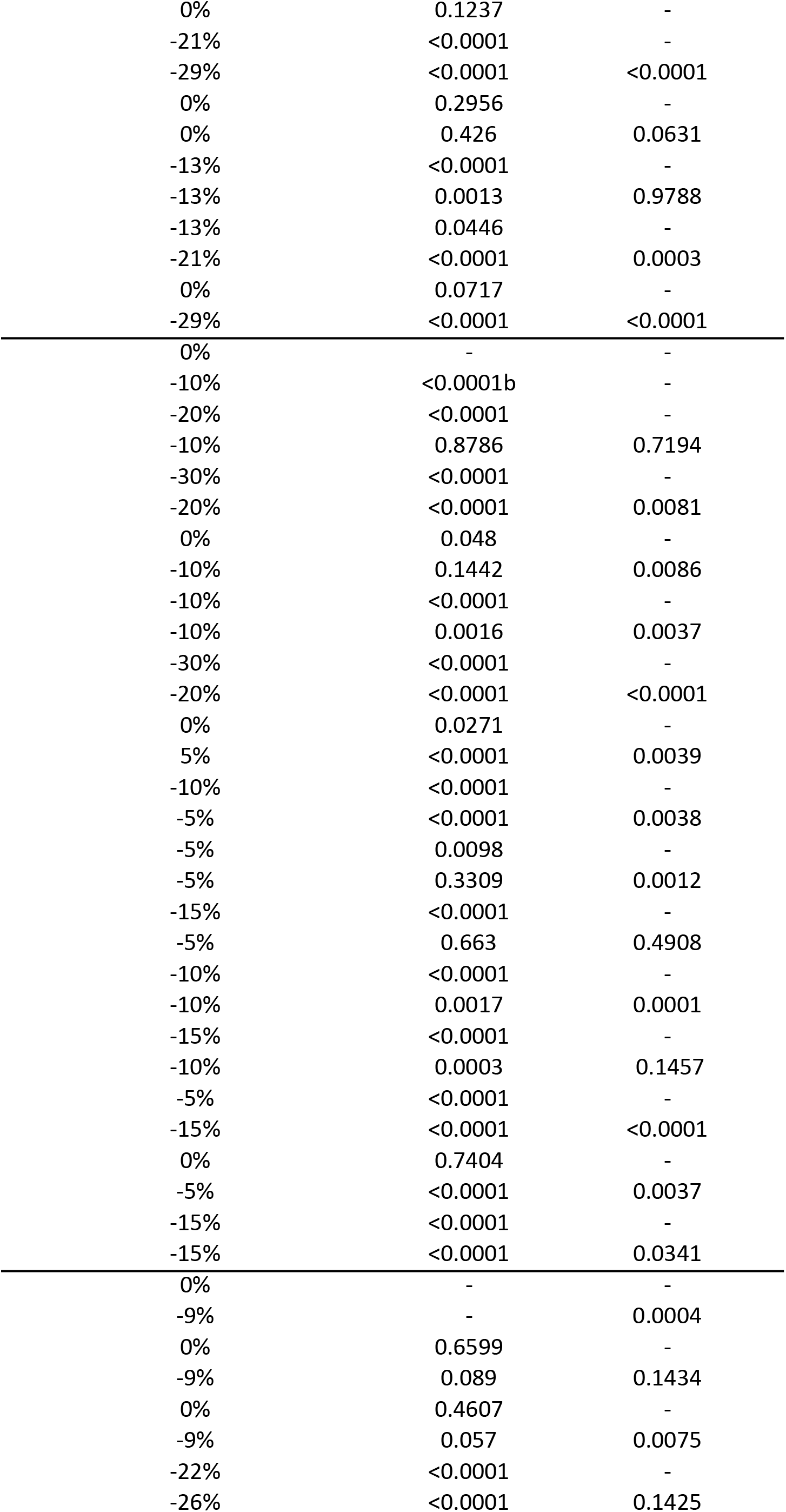
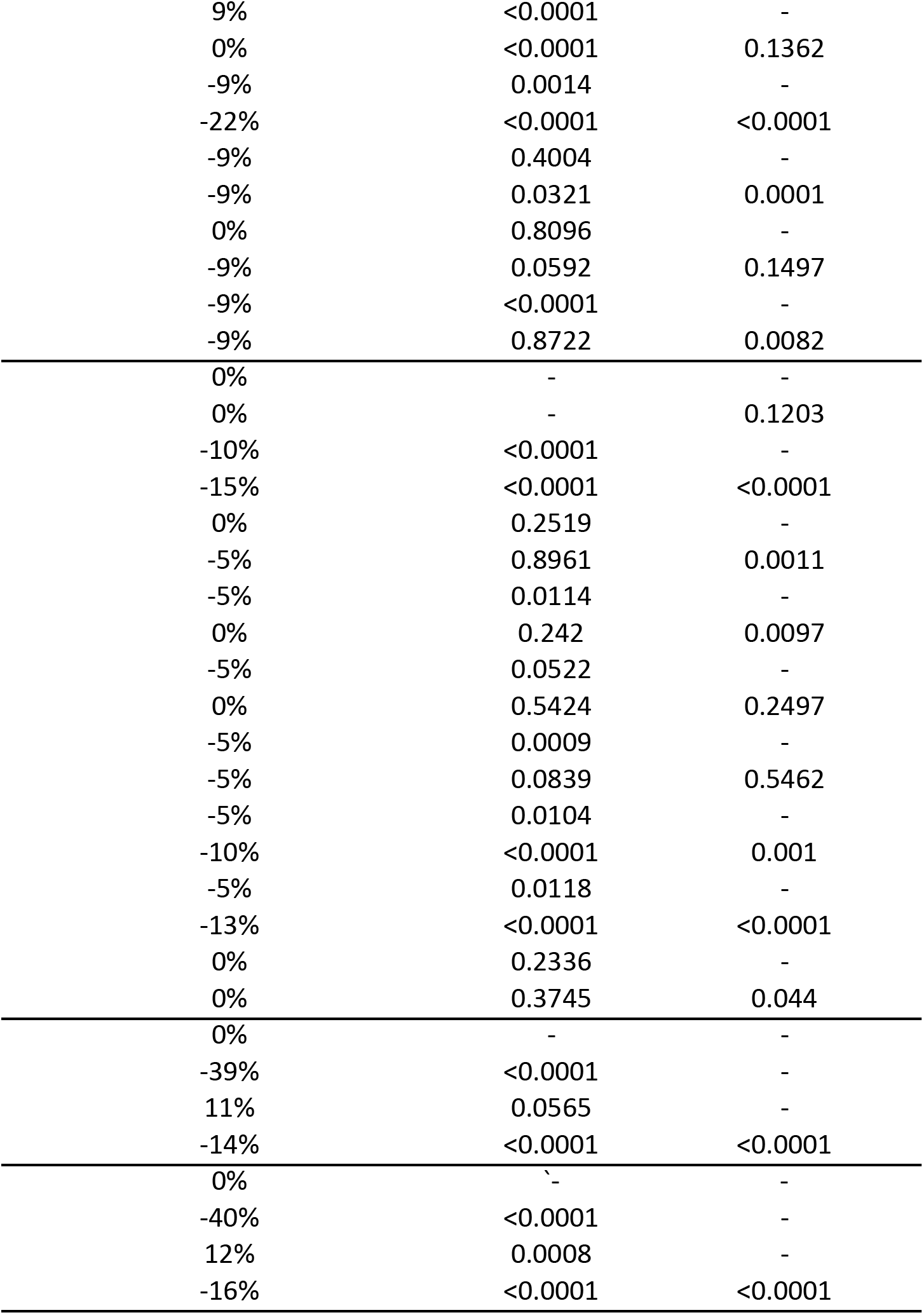

